# Genome-wide association study in Collaborative Cross mice reveals a role for *Rhbdf2* in skeletal homeostasis

**DOI:** 10.1101/094698

**Authors:** Roei Levy, Clemence Levet, Keren Cohen, Matthew Freeman, Richard Mott, Fuad Iraqi, Yankel Gabet

## Abstract

Osteoporosis, the most common bone disease, is characterized by a low bone mass and increased risk of fractures. Importantly, individuals with the same bone mineral density (BMD), as measured on two dimensional (2D) radiographs, have different risks for fracture, suggesting that microstructural architecture is an important determinant of skeletal strength. Here we took advantage of the rich phenotypic and genetic diversity of the Collaborative Cross (CC) mice. Using microcomputed tomography, we examined key structural parameters in the femoral cortical and trabecular compartments of male and female mice from 34 CC lines. These traits included the trabecular bone volume fraction, number, thickness, connectivity, and spacing, as well as structural morphometric index. In the mid-diaphyseal cortex, we recorded cortical thickness and volumetric BMD.

The broad-sense heritability of these traits ranged between 50 to 60%. We conducted a genome-wide association study to unravel 5 quantitative trait loci (QTL) significantly associated with 6 of the traits. We refined each locus by combining information obtained from the known ancestry of the mice and RNA-Seq data from publicly available sources, to shortlist potential candidate genes. We found strong evidence for new candidate genes, including *Rhbdf2*, which association to trabecular bone volume fraction and number was strongly suggested by our analyses. We then examined knockout mice, and validated the causal action of *Rhbdf2* on bone mass accrual and microarchitecture.

Our approach revealed new genome-wide QTLs and a series of genes that have never been associated with bone microarchitecture. This study demonstrates for the first time the skeletal role of *Rhbdf2* on the physiological remodeling of both the cortical and trabecular bone. This newly assigned function for *Rhbdf2* can prove useful in deciphering the predisposing factors of osteoporosis and propose new investigative avenues toward targeted therapeutic solutions.

**Author summary:** In this study, we used the novel mouse reference population, the Collaborative Cross (CC), to identify new causal genes in the regulation of bone microarchitecture, a critical determinant of bone strength. This approach provides a clear advantage in terms of resolution and dimensionality of the morphometric features (versus humans) and rich allelic diversity (versus classical mouse populations), over current practices of bone-related genome-wide association studies.

Our genome-wide study revealed 5 loci significantly associated with microstructural traits in the cortical and trabecular bone. We found strong evidence for new candidate genes, in particular, *Rhbdf2.* We then validated the specific role of *Rhbdf2* on bone mass accrual and microarchitecture using knockout mice. Importantly, this study is the first demonstration of a physiological role for *Rhbdf2*.

## Introduction

Osteoporosis is the most common bone disease in humans, affecting nearly half the US and European population over the age of 50 years. With the globally increasing life expectancy, osteoporosis and related bone fractures are becoming a pandemic health and economic concern. By 2050, the world-wide incidence of hip fractures is expected to increase by 2.5 to 6 fold [1,2]. Importantly, the mortality rate in the 12 months following bone fracture is as high as 20% [3]. Risk of fracture is determined largely by bone density and quality/strength, which are the end result of peak values achieved at skeletal maturity and subsequent age and menopause-related bone loss. Genetic factors have a major role in determining the wide range in the so-called “normal” peak bone mass. Measures of bone status are inherently complex traits, as opposed to Mendelian traits; i.e. they are controlled by the cumulative effect and interactions of numerous genetic loci and environmental factors.

Genome-wide association studies (GWAS), including a large meta-analysis, have identified more than 50 loci associated with bone mineral density (BMD) [4–10]. However, many other genes that were experimentally associated with bone mass were not confirmed by GWAS in human cohorts [11–13]. This suggests that the BMD phenotype does not capture the structural complexity of the bone; there may be other relevant bone phenotypes not yet studied in human GWAS [12], which hitherto have generally relied on areal bone mineral density (aBMD) as the sole bone feature. aBMD measured by dual energy x-ray absorptiometry (DXA) is a two-dimensional projection that cannot measure bone size, individual bone compartments’ shape (whether trabecular or cortical) or underlying microstructure, and thus likely conceals important features which are assumed to be controlled by unique genetic determinants. Indeed, there is a growing body of evidence that argues for distinct genetic influences of the cortical and trabecular bone and thus they should be accordingly distinguished [8,9]. A recent GWAS in the Collaborative Cross (CC) mice based on DXA failed to find any heritability of BMD [14], whereas another report based on the same mouse panel showed highly significant heritability levels in most of the cortical and trabecular microstructural parameters measured by micro-computed tomography (μCT)[15]

Traditional peripheral quantitative CT (pQCT) has the capacity to distinguish between the cortical and trabecular bone compartments, but it lacks the required resolution to detect microstructural differences. A recent report based on high resolution pQCT (HR-pQCT) data in humans, identified two novel bone-related loci, thus far undetected by DXA and pQCT-based GWAS [9], Another [16], found strong genetic correlations between 1047 adult participants of the Framingham heart study, therefore indicating that the heritability of bone microstructure constitutes a phenotypic layer which is at least partially independent of DXA-derived BMD. Like HR-pQCT studies in humans, understanding the genetic regulation of bone microstructural parameters using μCT in small animals is likely to identify genetic factors distinct from those previously identified for DXA-derived traits.

The CC mouse panel is designed to provide high resolution analysis of complex traits, with particular emphasis on traits relevant to human health [17,18], This unique resource currently consists of a growing number of recombinant inbred lines (RIL) generated from full reciprocal breeding of eight divergent strains of mice [19]. In contrast to commonly used laboratory mouse strains, the ancestry of the CC lines includes wild-derived inbred strains that encompass genetic variations accumulated over ~1 million years [20]; more than 50 million single nucleotide polymorphisms segregate in founders of the CC. The high genetic diversity means that QTLs can be mapped using this panel that would have been invisible in a population that involved only classical strains [21,22].

This claim is substantiated in a recent study that identified a genome-wide significant association between *Oxt* (oxytocin) and *Avp* (vasopressin) and skeletal microarchitecture in CC mice [15]. Here, our GWAS in the CC mouse panel identified a novel gene, *Rhbdf2*, associated with bone traits and using a specific knockout model we validated its role in the regulation of cortical and trabecular bone structure. This exemplifies the effectiveness and relative ease by which a GWAS with a small CC population can associate a bone-related function to novel genes, and to reveal overlooked key players in skeletal biology.

## Results

### CC lines widely differ in bone microarchitecture traits

We examined the variation in femoral cortical and trabecular microstructure between 34 unique CC lines totaling in 174 mice (71 females and 103 males, with an average of 4.25 mice per line). In the trabecular bone compartment we measured bone volume fraction (BV/TV), trabecular number (Tb.N), thickness (Tb.Th), connectivity (Conn.D), and spacing (or separation; Tb.Sp), as well as structural morphometric index (SMI) of the trabecular framework. In the mid-diaphyseal cortex, we recorded cortical thickness (Ct.Th) and volumetric bone mineral density (vBMD). These traits were approximately normally distributed; BV/TV ranged from 0.017 to 0.26 (i.e. 1.7% to 26%; mean = 10.2%); Tb.N from 0.52 to 6.11 mm^−1^ (mean = 2.7 mm^−1^); Tb.Th from 31 to 69 μm (mean = 47 μm); Conn.D from 10.9 to 268.3 mm^−3^ (mean = 104.2 mm^−3^); SMI from 0.6 to 3.3 (mean = 2.3); Tb.Sp from 0.16 to 0.7 mm (mean = 0.33 mm); Ct.Th from 0.14 to 0.29 mm (mean = 0.2 mm); and vBMD from 402.5 to 809.2 mgHA/cm^3^ (mean = 581.1 mgHA/cm^3^). μCT images taken from two mice with distinct cortical (Fig. 1A) and trabecular (Fig. 1B) characteristics demonstrate the great variation in bone traits due solely to the genetic background. Color-codes on the graphs in Fig. 2 indicate Duncan’s least significance range (LSR), which dictates whether the mean value of a line, or a group of lines, for a given trait differs to a degree of at least P-value < 0.001 from any other group. LSR allows for a visual representation of the heterogeneity amongst the lines.

**Figure 1.**
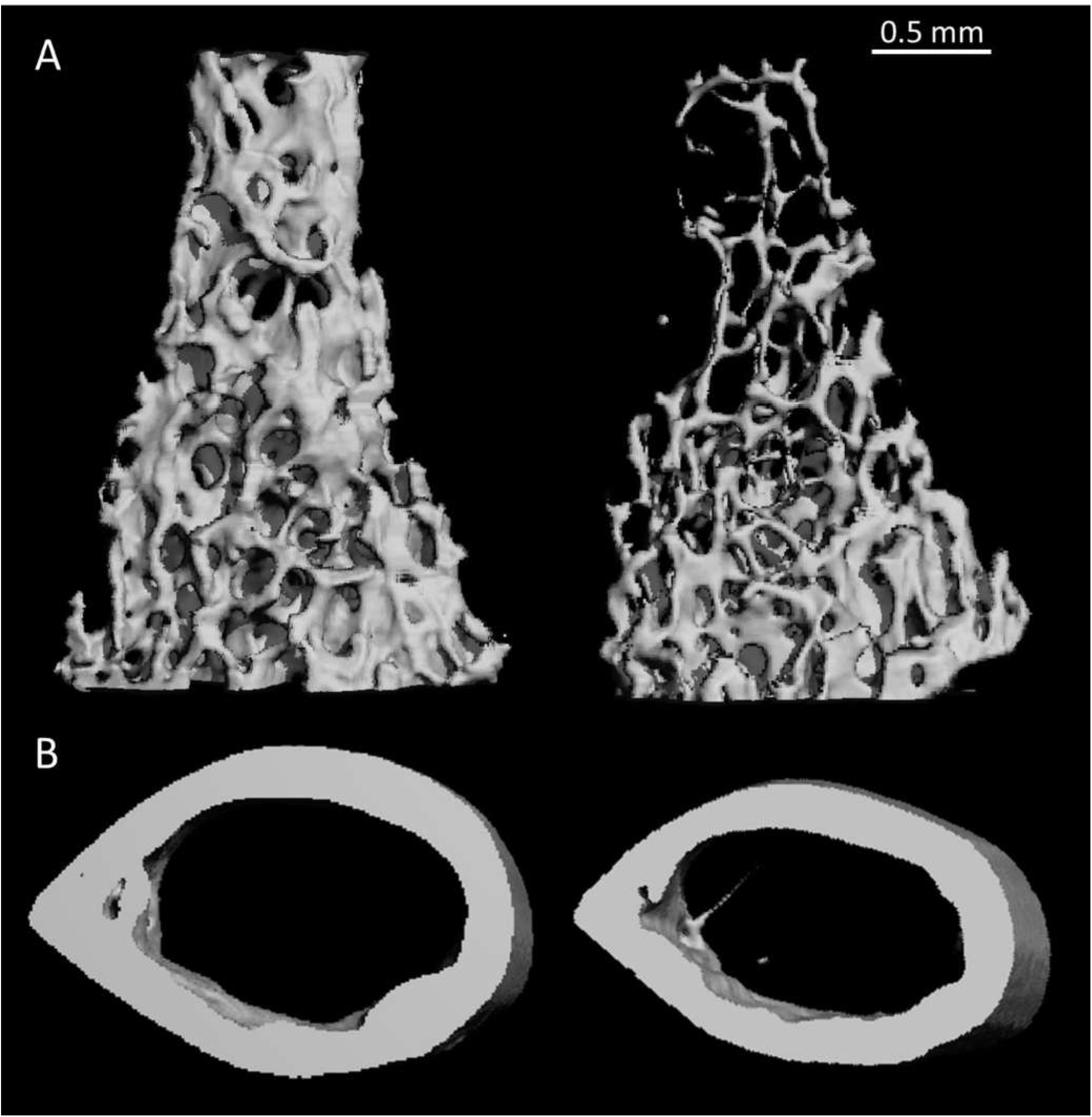
μCT images of trabecular and cortical bone of the femora of representative male CC mice. **(A)** Trabecular bone. Left: IL-2452, Right: IL-1513. **(B)** Cortical bone. Left: IL-785, Right: IL-2689.

**Figure 2.**
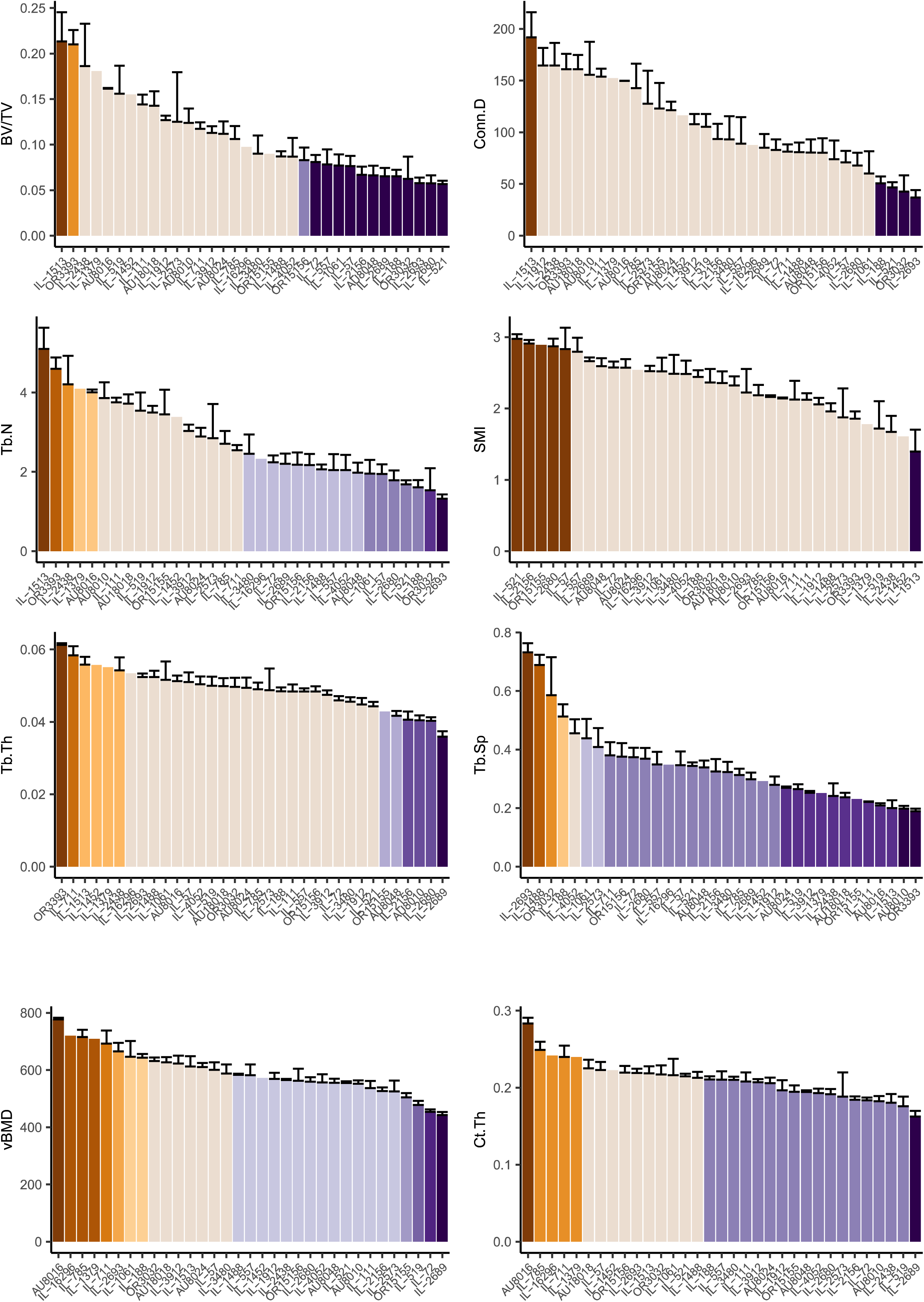
Trabecular and cortical traits distributions across the CC lines. X-axis is the lines, y-axis is the trait means. **A.** From top left, counter-clockwise: BV/TV (%), Tb.N (mm^-1^), Tb.Th (um), Conn.D (mm^-3^), SMI, and Tb.Sp (mm). **B**. Left, vBMD (mgHA/cm^3^), right, Ct.Th (mm). Color codes group line(s) which significantly differ from other groups. Lines are ordered inconsistently among the traits, per trait-specific descending order. Refer to Table S3 for more details.

With 11 distinct groups, vBMD (Fig. 2B) is the most heterogeneous trait, while SMI and Conn.D are the least, with only 3 significantly distinct groups (Fig. 2A). Notably, the heterogeneity of females is greater than that of males for cortical traits but milder for trabecular traits (Figs. S1A1, 2 for males and S1B1, 2 for females).

To examine the inter-dependency between the traits, we assessed the correlation between all the measured parameters, in a pairwise fashion using Pearson’s correlation test. The strongest correlation was between BV/TV and Tb.N (Pearson’s *r* = 0.94), in line with our previous findings [15], while the weakest was between Tb.N and vBMD *(r* < 0.01). There was also a moderately high correlation between Ct.Th and Tb.Th *(r =* 0.61; and see Table S1). The correlation between sexes for each trait (Table 1) ranged from *r* = 0.75 (Tb.Sp) to *r* = 0.20 (Ct.Th). Body weight (range = 17.4 - 35.0 gr) did not significantly correlate with any of the traits (*r* = 0.01 for Conn.D to *r* = 0.19 for Ct.Th). After separating males from females the correlation slightly increased, yet remained low. Weak correlation was found between weight and Tb.N, SMI, and Ct.Th for females (Pearson’s *r* = −0.20, 0.23, and 0.25 respectively), and between weight and Tb.Th and Tb.Sp (Pearson’s *r* = 0.25 and −0.25) for males.

**Table 1.** Heritability, sex correlations, and covariate interactions for trabecular and cortical traits

| **Trait** | **H<sup>2</sup>** | **logP** | **H<sup>2</sup>n** | **Sex Cor.** | **Interactions %** |
| --- | --- | --- | --- | --- | --- |
| BV/TV | 0.61 | 12.43 | 0.87 | 0.6695754 | - |
| Tb.N | 0.63 | 13.76 | 0.88 | 0.7628464 | - |
| Tb.Th | 0.54 | 9.08 | 0.83 | 0.7039887 | 34.70 |
| Conn.D | 0.56 | 9.90 | 0.84 | 0.5188039 | - |
| SMI | 0.55 | 9.43 | 0.84 | 0.2984166 | 26.02 |
| Tb.Sp | 0.63 | 13.45 | 0.88 | 0.754563 | - |
| vBMD | 0.62 | 12.15 | 0.87 | 0.6268818 | 53.92 |
| Ct.Th | 0.51 | 7.63 | 0.82 | 0.2055684 | 41.07 |

H^2^ is the broad-sense heritability (which includes epistatic and environmental influences); logP is the negative 10-base logarithm of the P-value; H^2^n is the line-mean heritability; Sex Cor. Is the sex correlation of each trait; and interactions % refers to the relative contribution of the cumulative covariate-interactions, which include sex, age, batch, month, season, year, and experimenter (see table S2).

While in most lines the traits’ correlations were predictive of a given line’s rank, in others a less expected pattern was observed; e.g., IL-1513 displayed unusually extreme phenotypes for all trabecular traits and was at the higher end for BV/TV, Tb.N, Tb.Th, and Conn.D and at the lower end for SMI and Tb.Sp, but IL-188 was more discordant between these same traits (Fig. 2 and Table S3), illustrating unexpected covariation of the traits in the CC.

### Heritability and confounder-control

We quantitated the effects of the covariates sex, age, batch, month, season, year, and experimenter on each trait. Age ranged from 9 (n=6) to 13 (n=9) weeks and the mice were dissected in 20 batches over a three-year course across 8 months during winter, spring and summer, by two experimenters. Whereas age alone had no effect on any trait, sex affected only Ct.Th; batch affected Tb.Th, vBMD, and Ct.Th; month affected Tb.Sp, vBMD, and Ct.Th; season and year affected vBMD and Ct.Th; and Tb.Th and Ct.Th were affected by experimenter. The cumulative effect of the covariates’ pairwise interactions was noted for Tb.Th, SMI, vBMD, and Ct.Th. (Table S2).

We then estimated the broad-sense heritability (H^2^) of each trait among the CC lines, which includes additive and non-additive epistatic effects and gene-environment interactions. The greatest H^2^ is seen for Tb.N (0.63, logP = 13.76; where logP stands for the negative 10-base logarithm of the P value and tests the null hypothesis that the heritability is zero), and the smallest for Ct.Th (0.51, logP = 7.63).

We calculated the heritability for the mean values in each line to get a better representation of the percentage of genetic contribution to the phenotypic heterogeneity by incorporating H^2^ and the average number of lines [23]. This defines H^2^n, which is directly proportional to H^2^ (Methods; Table 1) and ranges between 82 (Ct.Th) and 88% (Tb.N and Tb.Sp).

Overall the cortical traits seemed more prone to covariate variation; they were particularly sensitive to sex, batch, and season. This stands in contrast to our previous results [15] where BV/TV, Tb.N, and Conn.D displayed a profound sex effect, although there cortical traits were not measured. This means there is a deeper, complex layer of sex effect dependent upon cooperative environmental and genetical factors which requires further work to fully comprehend.

### Association analysis for microarchitectural traits highlights 5 QTLs

We first measured statistical association between each trait and the founder haplotype at each locus in the genome. Association analyses of the cortical and trabecular traits to the haplotypes segregating in the CC (as defined by the ~70 K MegaMuga SNPs) yielded 5 distinct QTLs. For BV/TV and Tb.N we recognized a marked peak at a locus of length ~0.45 Mb between 116.5 and 116.9 Mb on chromosome 11, with peak logP values of 7.6 and 6.8, which extended above the 99^th^ percentile permutation-threshold by 2.7 and 1.94 logP units, respectively. In Tb.Th, Tb.Sp, Ct.Th, and vBMD we identified different QTLs on chromosomes 4, 5, 4 and 3, with logPs of 8.0, 9.4, 8.2, and 9.8, respectively, above threshold (Fig. 3 and Table 2). To account for false positive results, we kept the false discovery rate (FDR) at 1% for each scan, by employing the Benjamini Hochberg multiple testing procedure on the logPs of the haplotype associations to the traits. Conn.D and SMI lacked significant peaks above the stringent permutated threshold and thus were not further analyzed (Fig. S2), but Conn.D displayed a borderline peak in a region that matches the peak identified for BV/TV and Tb.N. The 5 QTLs we describe are hereafter referred to as *Trl* (trabecular related locus) 7-9, and *Crl* (cortical related locus) 1-2 (respectively for BV/TV and Tb.N, Tb.Th, Tb.Sp, Ct.Th, and vBMD, and in keeping with our previous report [15] that introduced *Trl* 1-6). The 95% widths of the confidence intervals ranged from 6.4 to 15.6 Mb for the *Trl*s, and were between 8.5-10.8 Mb for the *Crl*s (Table 2 and Fig. S3).

**Table 2.** Positions of QTLs associated with trabecular and cortical traits.

|  |  |  |  |  |  |  |  |  |  |  |  |
| --- | --- | --- | --- | --- | --- | --- | --- | --- | --- | --- | --- |
| QTL | Trait | Chr | logP | 99th % thresh | H<sup>2</sup><sub>r</sub> | Simulation |  |  |  |  |  |
|  |  |  |  |  |  | 50% CI (Mb) |  | 90% CI (Mb) |  | 95% CI (Mb) |  |
|  |  |  |  |  |  | Position | Width | Position | Width | Position | Width |
| *Trl7* | BV/TV | 11 | 7.60 | 4.90 | 0.69 | 116.6-116.7 | 0.12 | 113.6-118.1 | 4.50 | 112.1-118.3 | 6.40 |
| *Trl7* | Tb.N | 11 | 6.80 | 4.84 | 0.71 | 116.6-116.87 | 0.29 | 114.2-118.35 | 4.15 | 112.41-118.68 | 6.27 |
| *Trl8* | Tb.Th | 4 | 8.00 | 4.50 | 0.61 | 117.2-117.58 | 0.32 | 113.05-125.54 | 12.49 | 110.87-126.52 | 15.65 |
| *Trl9* | Tb.Sp | 5 | 9.40 | 6.01 | 0.78 | 105.78-106.14 | 0.35 | 101.6-109.11 | 7.54 | 99.8-110.39 | 10.62 |
| *Crl1* | Ct.Th | 4 | 8.20 | 4.80 | 0.80 | 9.29-9.72 | 0.43 | 4.0-11.7 | 7.70 | 3.4-11.8 | 8.49 |
| *Crl2* | vBMD | 3 | 9.80 | 4.93 | 0.86 | 97.2-97.4 | 0.20 | 94.4-103.1 | 8.50 | 93.22-104.3 | 10.80 |
| Chr = chromosome; logP = negative 10-base logarithm of P value; Sig = genome-wide significance level; 99th % threshold logP = threshold used to define cut-off for QTL peaks (Fig. 4); H<sup>2</sup><sub>r</sub> = regional heritability (the proportion to which the locus explains the phenotypic variability). Positions and widths of the simulation-based 50, 90, and 95% CIs are given. |  |  |  |  |  |  |  |  |  |  |  |

**Figure 3.**
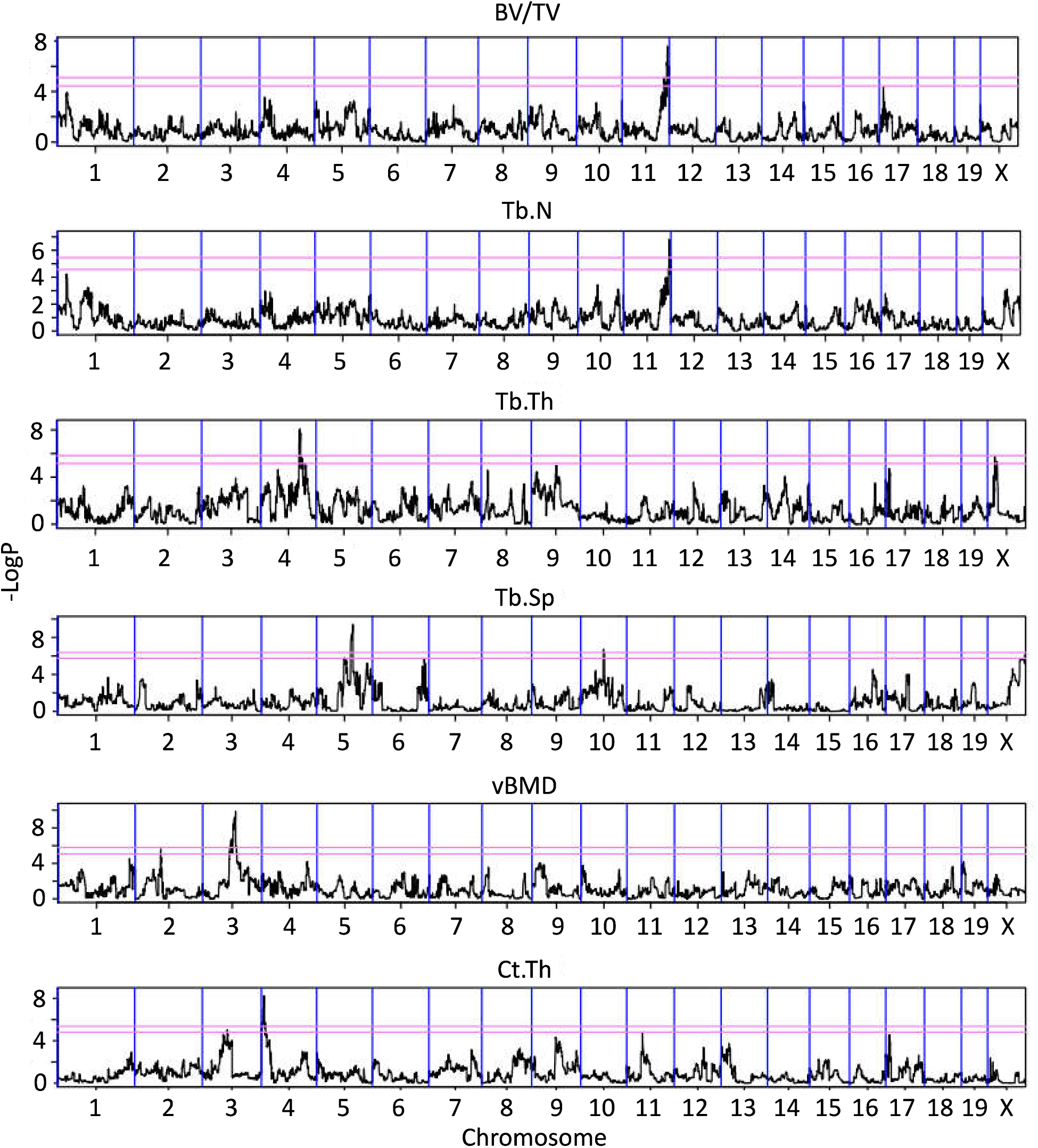
Haplotype association maps for the trabecular and cortical traits. X-axis is the position on the chromosome, y-axis is the −logP value of the association. Lower threshold represents the 95^th^ percentile of 200 simulations, and top represents the 99^th^ percentile. Loci above the 99% cut-off were further investigated. From top to bottom: BV/TV, Tb.N, Tb.Th, Tb.Sp, vBMD, and Ct.Th.

We measured the contribution of each CC founder to the QTLs, relative to the wild-derived strain WSB/EiJ (Fig. 4). *Trl*7 is mostly affected by the classic laboratory strains 129S1/SvImJ, NOD/LtJ, and NZO/HiLtJ; notably, the other traits were more strongly driven by the following wild strains: *Trl*8 and *Crl*2 by PWK/PhJ; *Trl*9 by WSB/EiJ; and *Crl*1 by CAST/Ei.

**Figure 4.**
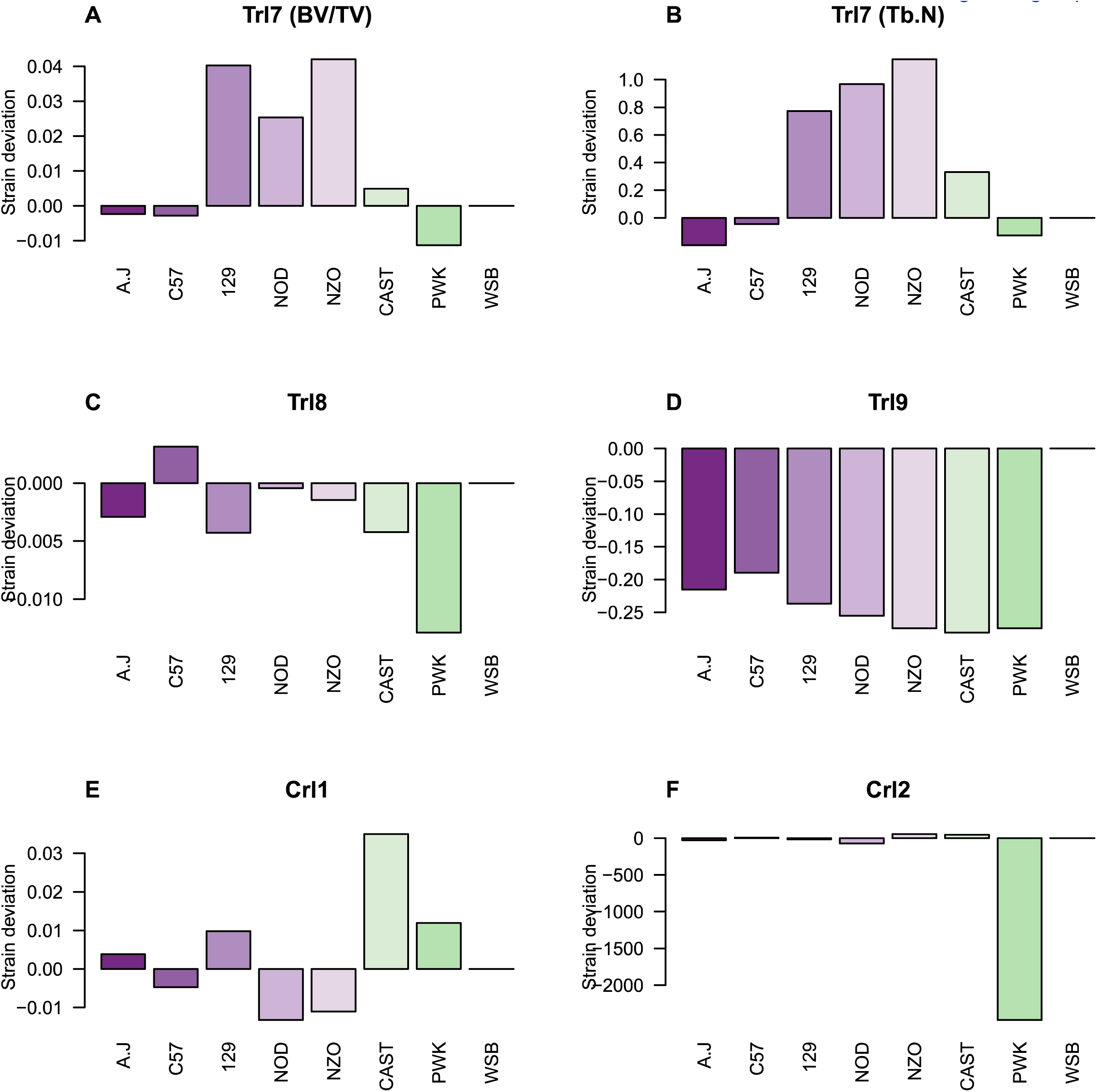
Ancestral effects relative to WSB. Y axis is the strain deviation relative to WSB, x axis is the different strains of the eight CC founders. **(A)** to **(F)**: *Trl7* to *Trl9, Crll*, and *Crl2*, respectively.

For *Trl7*, at the SNP most adjacent to the QTL peak UNC20471277, we found that the majority of lines with a TT allelic variant (where T refers to the nucleic acid Thymine) mostly congregate at the higher end of the BV/TV and Tb.N values (Mean BV/TV = 17%); lines with a CC variant (where C refers to the nucleic acid Cytosine) are at the lower end (mean BV/TV = 10%); and those with a CT variant are at the intermediate range (Fig. 5). Largely, the more the trait examined is distantly correlated with BV/TV, the less differentiated the CC and TT variants are, at the SNP UNC20471277. This is accentuated in vBMD where there is a weak correlation with BV/TV (Table S1) and leveled CC and TT groups (*P* value = 0.8 Welch’s two sample t-test).

**Figure 5.**
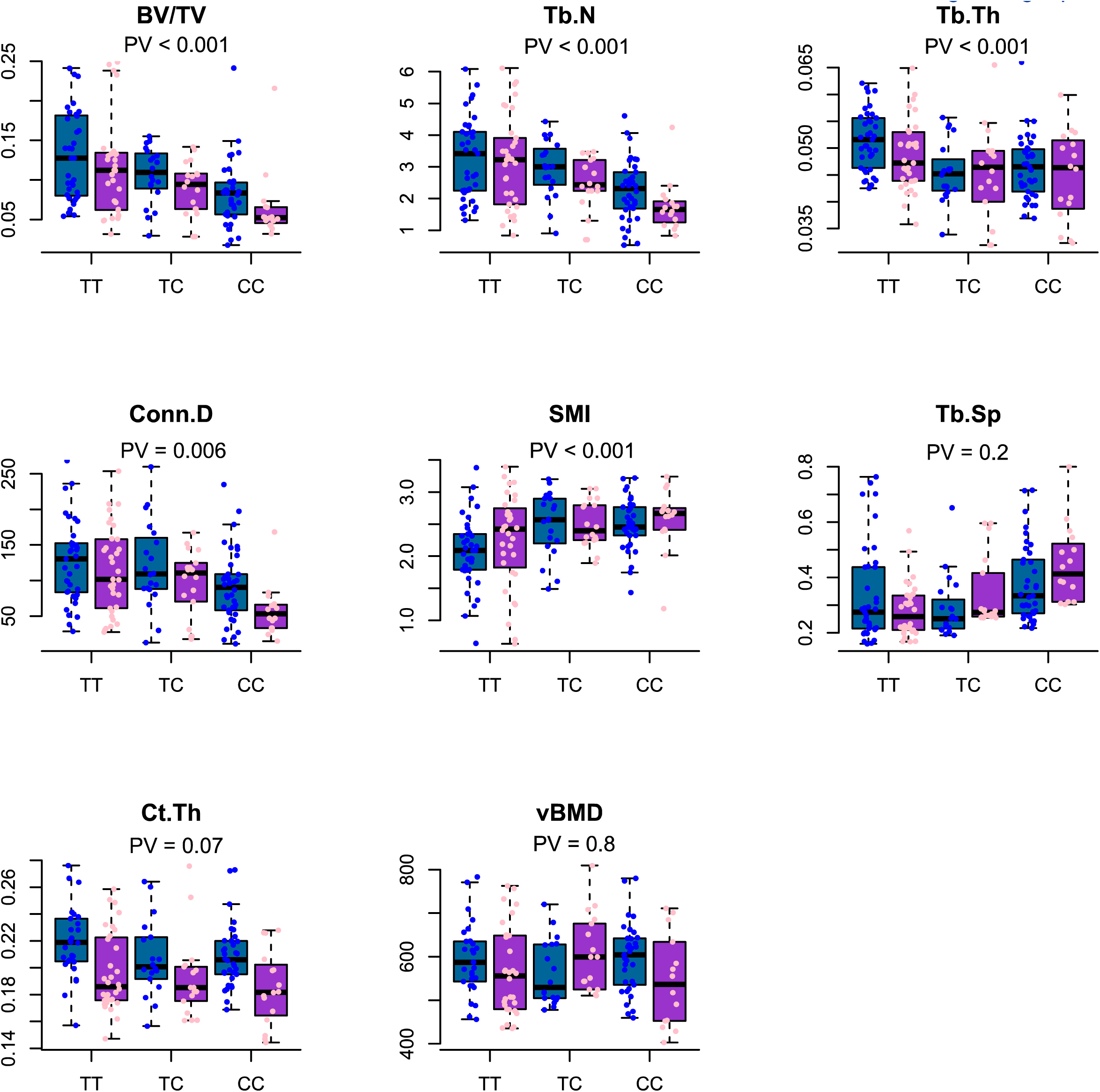
Traits distribution at the marker UNC20471277 across bearers of homozygous and heterozygous alleles, separated by sex. X-axis is the allelic variation at the marker, y-axis is the trait value.

**Figure 6.**
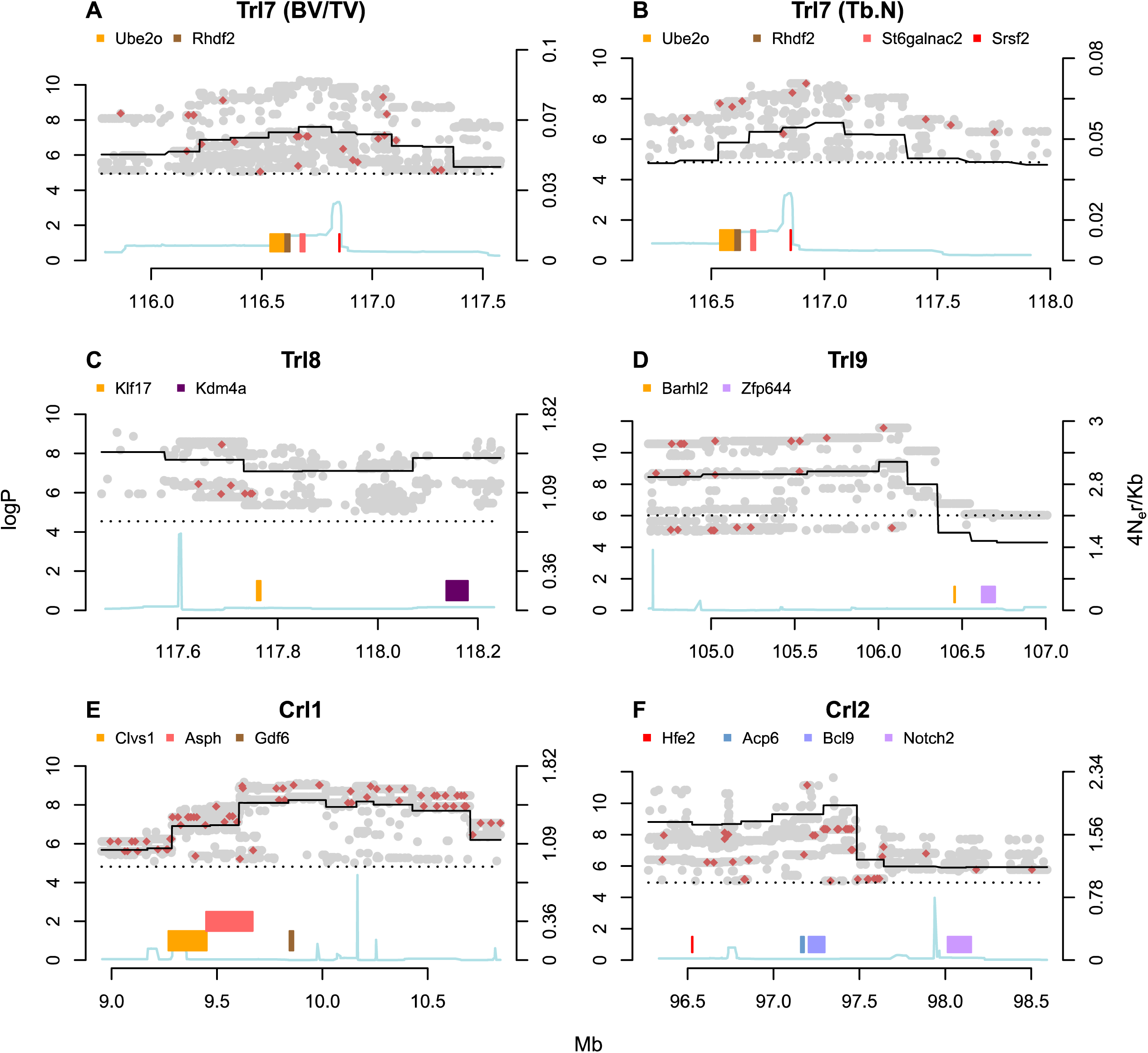
Merge analysis. Readings below logP = 4 are elided for brevity. X axis is the position on the genome in Mb; y left axis is the logP score; y right axis is the recombination rate scale; colored bars are genes (note that only strong putative candidate genes are shown.); cyan line is the recombination rate; black continues line is the haplotype test’s peak; dashed line is the 99% permutation threshold. **(A)** to **(F)**: Trl7 of BV/TV, Trl7 of Tb.N, Trl8 of Tb.Th, Trl9 of Tb.Sp, Crl1 of Ct.Th, and Crl2 of vBMD, respectively.

### Candidate genes identified by merge analysis and RNA-seq

To identify the gene most likely driving the skeletal trait, we next performed a merge analysis and RNA-seq analysis and used a scoring system to rank the potential candidates.

Merge analysis uses the catalogue of variants segregating in the eight CC founders to impute the genotype dosage of each SNP in each CC line, based on the haplotype reconstruction used for haplotype association [24]. Candidate causal variants, if they exist, would be expected to be more significant (have higher logP values) than the haplotype-based test in the flanking region. We found that *Trl7* had the highest density of polymorphisms (grey and crimson dots in Fig. 6) with merge-logP values above the haplotype logPs (continuous black line in Fig. 6), while *Trl*8 and *Crl*2 had very few. The merge logP values of the two latter loci congregated more upstream, in accordance with the left-skewness of their respective CI simulations (Fig. 6; Fig. S3). By calculating the relative density of merge logP values which are considerably higher than the haplotype merge logPs -and above the 99^th^ percentile threshold for each scan - at intervals defined by each gene within the QTLs (in meaningful regions; usually between the 50th and 90th CI percentile) we could rank the genes according to their merge analysis results (Table S4). For example, while the proportion of merge analysis SNPs for BV/TV and Tb.N with logPs greater than that of the haplotype scan is 1.4% at the genome-wide scale (as well as at the region spanning the 95% CI, between ~112 – 118 Mb), it is 9.4% and ranked 5/36 (for BV/TV) or 14.07% and ranked 3/36 (for Tb.N) at the region in which the gene *Rhbdf2* is situated (~116.5 – 116.6 Mb) (See Table S4 and further discussion below).

To strengthen the criteria that classifies potential putative candidate gene as true positives, we analyzed RNA-seq datasets of osteoclasts (Fig. S4) and osteocytes (Fig. S5) made publicly available by Kim *et al* [25] (Gene Expression Omnibus accession number GSE72846) and St John *et al* [26] (Gene Expression Omnibus accession number GSE54784). We focused on local maximas that span ~0.5 Mb in and around the peaks suggested by the merge analysis, for each QTL. From the raw count reads we found that *Trl7* had the strongest gene expression differential; e.g while *Mxra7* in the osteoclasts was expressed to a negligible degree (Fig. S4A), it had a strong presence in osteocytes (Fig. S5A), whereas the genes of the other loci had much less prominent differences. This suggests that genes at Trl7 are differentially expressed between osteocytes and osteoclasts more prominently than in the other loci.

For each of the genes with the highest merge analysis scores in *Trl7*, we attributed an *osteoclast* and *osteocyte* RNA-seq (S_RNA-seq1_ and S_RNA-seq2_, respectively) score based on the following formula: 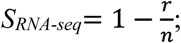 where *r* is the local gene rank (sorted by the raw read count) and *n* is the total number of genes at the locus, where n=10, if n>10. For example, *Ube2o* had an *osteoclast* RNA-seq score of 0.8 (ranked 2^nd^ out of n>10 genes) and *osteocyte* RNA-seq score of 0.8. *Rhbdf2* had an *osteoclast* RNA-seq score of 0.9 (ranked 1^st^) and osteocyte RNA-seq score of 0.7.

We then summed up the cumulative MS (*M*erge and *S*equencing) score for each gene, defined as *MS(i)* = *ln(Mstrength(i)*) + (*S_RNA-seq1_(i)* + *S_RNA-seq2_(i)*), where *M_strength_*, *S_RNA-seq1_*, and *S_RNA-seq2_* refer to the merge rank (or strength), osteoclast RNA-seq score, and osteocyte RNA-seq score of a given gene (*i*), respectively. Our analytical approach and scoring system enabled us to shortlist the most plausible causal genes at the QTLs. In *Tlr7, Ube2o* had an *MS_Ube2o_* = 2.4 + (0.8 + 0.8) = 4.0; while Aanat scored *MS_Aanat_* = 2.7 + (0 + 0) = 2.7, and *Rhbdf2* scored *MS_Rhbdf2_* = 2.6 + (0.9 + 0.7) = 4.2. In the other loci, the highest ranked genes were *Klf17* and *Kdm4a* for *Trl8*; *Barhl2* and *Zfp644* for *Trl9*; *Asph* and *Gdf6* for *Crl*1; and *Hfe2*, *Acp6*, *Bcl9*, and *Notch2* for *Crl*2.

Because BV/TV is a predominant parameter in bone biology, we first focused on *Trl*7. *Rhbdf2* had a merge strength of 14% (the 3^rd^ strongest at the QTL and 2^nd^ at the 50% CI), and a local maxima at the RNA-seq of the osteoclasts. It was located near the haplotype mapping peak, and because it received the highest cumulative *MS* score, *Rhbdf2* was retained for validation. The comprehensive list of the genes under the 50, 90, and 95% CI of the QTL, is supplied in table S4.

### Validation of the skeletal role of *Rhbdf2* in knock-out mice

Femora of male mice (n=14) null at *Rhbdf2* (on a C57B1/6J background) were collected, on which we measured the same morphometric traits as above, including BV/TV, Tb.N, Tb.Th, SMI, Tb.SP, Ct.Th, and vBMD. These were compared to their wild-type (WT) counterparts (n=13), after adjusting for batch, age and weight.

Strikingly, we found that *Rhbdf2*^-/-^ mice had a significant bone phenotype. In line with our GWAS data, *Rhbdf2*^-/-^ mice displayed a highly significant increase in BV/TV and Tb.N (Fig. 7, 8). As expected, *Rhbdf2* KO also affected other microstructural parameters, partly due to the high correlation between the trabecular traits. After adjusting for confounders, we observed a significant difference between KO and WT animals in Tb.Sp (P value < 0.001; uncorrected P value = 0.017), SMI (P value = 0.03; uncorrected P value = 0.156) and Conn.D (P value = 0.008; uncorrected P value = 0.046). Tb.Th and vBMD were not affected by the knockout. Although the cortical compartment did not display a haplotype peak at the vicinity of *Trl*7 in the CC animals. However, after adjusting for confounders, we observed a significant difference in Ct.Th between KO and WT bones (P value = 0.01), suggesting that the role of *Rhbdf2* is not limited to the trabecular compartment (Fig. 7).

**Figure 7.**
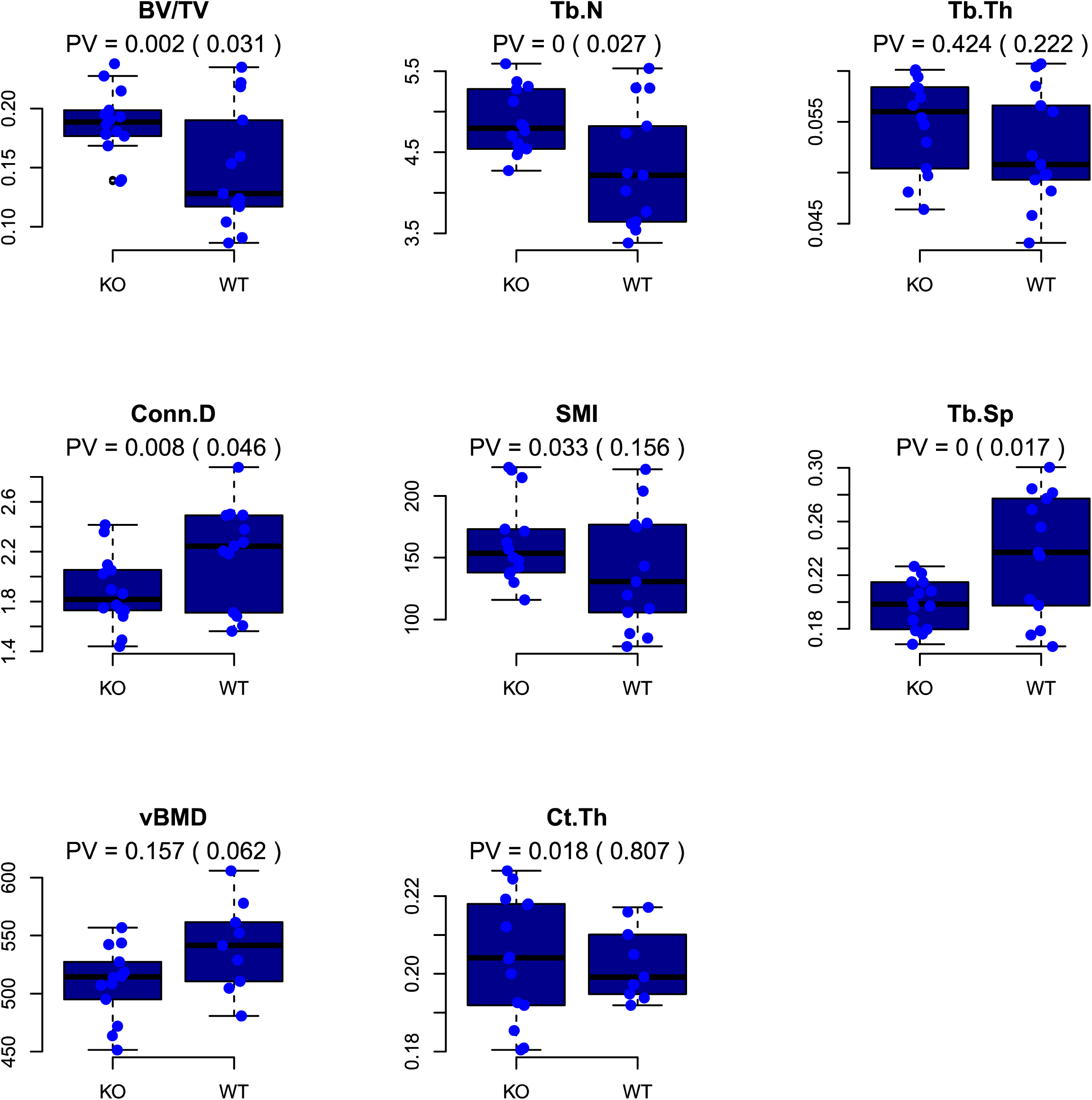
*Rhbdf2* knockout versus wildtype for each of the studied traits. Left is KO, right is WT. PV is the confounder-adjusted P value. The unadjusted P value is in brackets. PV = 0 means PV < 0.001.

**Figure 8.**
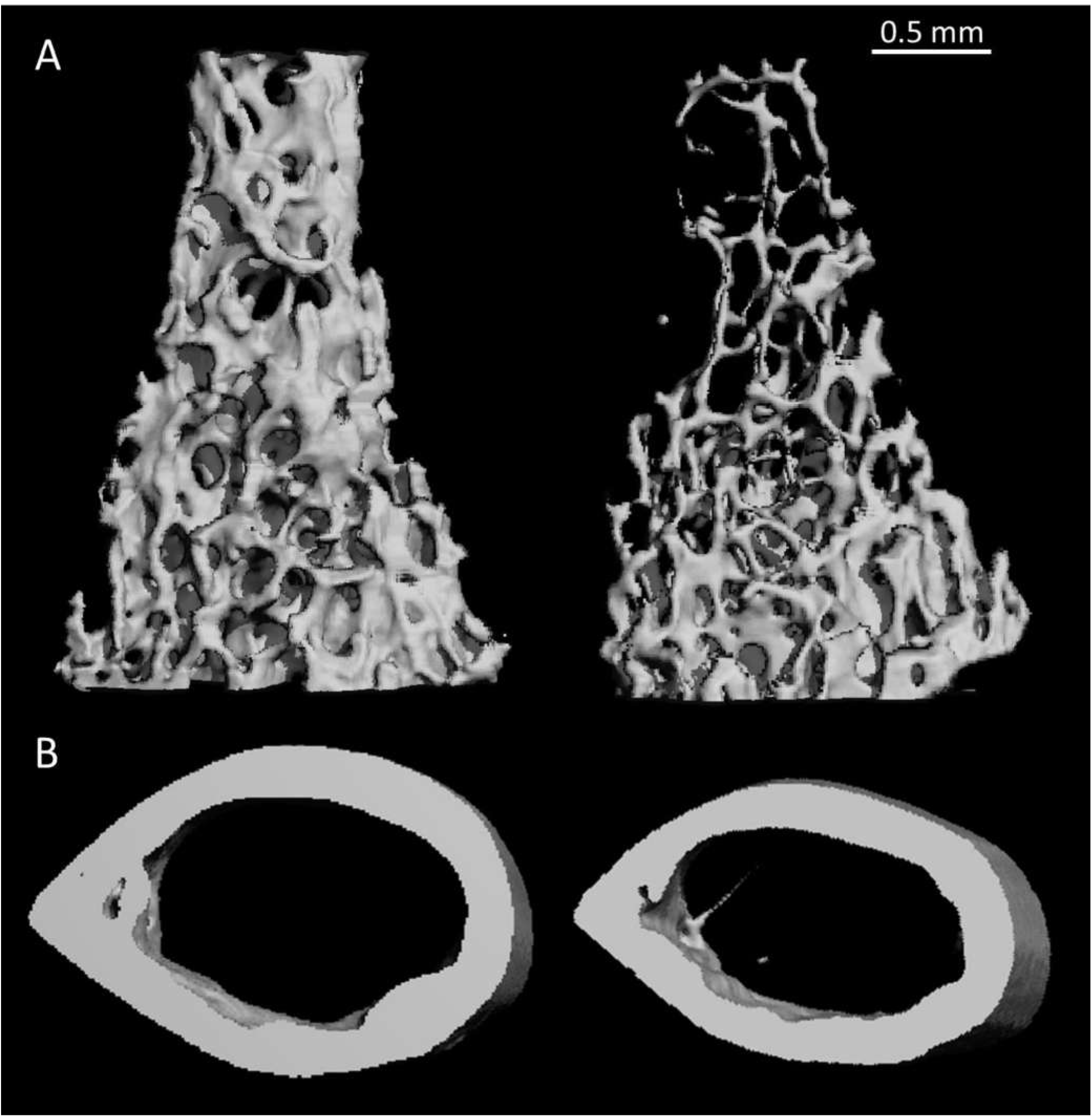
μCT images of three-dimensional representative cortical and trabecular bones reconstructions for *Rhbdf2* knockout and wildtype. Left: KO, right: WT. **(A)** Trabecular bone. **(B)** Cortical bone. All samples were of male mice, aged 11 weeks.

## Discussion

Genetic reference population (GRP) are very efficient for the study of complex traits and biological systems, because (i) genotyping is only required once (“genotype once, phenotype many times”, see below), and (ii) replicate individuals with the same genotype can be generated at will allowing for optimal experimental designs [27].

This article is the second to present the results of an ongoing quest to delineate the genetic determinants that govern microstructural bone traits [15]. Here we characterize several key microstructural properties of the mouse femoral bone to assess the extent to which they are heritable; to what environmental perturbations they are prone; and to identify candidate genes by which they are controlled. Our approach narrowed down a small number of putative candidate genes for 6 of the 8 examined phenotypes. Following merge analysis and RNA-seq, we validated our leading candidate gene, *Rhbdf2*, using a knockout model, which confirmed the critical role of *Rhbdf2* on bone mass accrual and homeostasis.

While the heritability rates assessed here - determined to be over 60% for all traits - confirmed our previous findings, the degree to which sex explains the phenotypic variation was very subtle, and appeared only for the cortical traits; this discrepancy may be due to the specific cohort composition used in this study (Table S3), which includes a sex bias due to smaller number of females than males. We found a total of five QTLs in six traits; BV/TV and Tb.N shared one QTL, and Tb.Th, Tb.Sp, vBMD, and Ct.Th yielded one each. Importantly, although bone microarchitecture factors are complex traits, our analyses highlighted no more than two loci for each trait; it is likely that analyzing a larger number of CC lines would result in the identification of further loci.

Our analyses yielded three QTLs for the trabecular traits and 2 QTLs for the cortical traits. These are referred to as *Trl*7-9, and *Crl*1-2, respectively. In addition, and in close proximity, to *Rhbdf2* (Rhomboid 5 Homolog 2; elaborated below), *Trl*7 includes *Ube2o* (Ubiquitin Conjugating Enzyme E2 O), which encodes an enzyme that is an important interactant of SMAD6. Ube2o monoubiquitinates SMAD6, and thereby facilitates the latter to bind BMP 1 receptors [28]. The signal transduction of BMP 1 is in turn limited [29,30], and endochondral bone formation, instead of ossification, is favored. Importantly, 4 week-old SMAD6-overexpressed mice have significantly lower humeral and vertebral BV/TV ratios than their controls [29].

At *Trl8*, *Klf1*7, *Kdm4a*, and *Dmap1* are likely putative candidate genes. Since *Klf1*7 (Kruppel-Like Factor 17) is part of a network that includes BMPs [31] it is more likely than a nearby gene, *St3gal3* (ST3 Beta-Galactoside Alpha-2,3-Sialyltransferase 3), to affect bone traits, although the latter has a greater merge strength. *Kdm4a* (Lysine Demethylase 4A) encodes a histone demethylase that promotes the differentiation of embryonal stem cells (ESCs) to an endothelial fate [32]; endothelial cells are implied in regulation of bone formation [33]. *Dmapl* (DNA Methyltransferase 1 Associated Protein) which encodes a DNA-methyl transferase known to regulate obesity complications, and is differentially methylated in women with polycystic ovary
syndrome [34,35] had the highest meaningful merge density, and it might epigenetically regulate bone formation as well. *Trl*9 includes two genes of interest to bone biology: *Barhl2* and *Zfp644*. By interacting with caspase3, which is essential for ossification [36], *Barhl2* (BarH Like Homeobox 2) can inhibit β-catenin activation [37], and regulate the expression of chordin, a BMP signaling-detrimental protein [38]. *Zfp644* (Zinc Finger Protein 644), which encodes a transcription repressor zinc-finger protein, is upregulated in eight week-old ovariectomized mice following treatment with estradiol [39], a treatment associated with reduced bone loss [40]. Further support for the candidacy of *Barhl2* and *Zfp644* is given by the role of *Barhl2* in the development of amacrine cells [41,42] and the association of *Zfp644* with myopia [43], a condition speculated to propagate from amacrine cell signaling [44]; interestingly myopia was linked to reduced postnatal bone mineral content in humans [45] and decreased expression of BMP 2 and 5 in guinea pigs [46].

The first of two cortical loci, *Crl*1 contains as likely candidates the genes *Asph* and *Gdf6. Asph* (Aspartate Beta-Hydroxylase) encodes a protein that has a role in regulating calcium homeostasis, which may affect bone metabolism [47]. *Gdf6* (Growth Differentiation Factor 6) is bone morphogenetic protein 13: mice with mutated *Gdf6* exhibit deformed bone formation in various skeletal sites; it is among the earliest known markers of limb joint formation [48], expressed in joints of ankle and knee. In *Gdf6* homozygous mutant mice, bones fuse at the joints early at the segmentation stage [49], For *Crl*2, we found *Hfe2, Bcl9, Notch2*, and *Prkab2* as potential candidate genes. *Hfe2* (Hemochromatosis Type 2 (Juvenile)) encodes the BMP co-receptor hemojuvelin which is expressed in skeletal muscles [50] and is responsible for juvenile hemochromatosis, a condition linked to sex hormones depletion and osteoporosis [51]. *Bcl9* (B-Cell CLL/Lymphoma 9), the mammalian ortholog of the gene *Legless*, encodes a protein essential to the Wnt/beta-catenin signaling which is important for bone metabolism [52], without which the nuclear localization of β-catenin and myocyte differentiation are compromised [53]. Of note, there are mutual effects between bone and muscle, and accumulating evidence suggest many genes show pleiotropism with respect to muscle strength and bone parameters [54], *Notch2* encodes a member of the notch protein family, which influence both osteoblasts and osteoclasts [55]; specifically, *Notch2* is associated with the rare Hajdu-Cheney syndrome, that includes severe osteoporosis as one of its main symptoms [56,57]. For this gene, we did not find any significant merge logPs included within its limits. Interestingly, *Sec22b*, an adjacent gene, had the strongest merge logP marks in this locus but no documented link to bone biology. The third-strong gene in terms of merge values was *Prkab2* (Protein Kinase AMP-Activated Non-Catalytic Subunit Beta 2). It encodes an enzyme which is the regulatory subunit of mitogen-activated protein kinase (AMPK). AMPK widely affects bone metabolism [58].

*Rhbdf2* is not yet supported by peer-reviewed reports as bearing a relation to bone. Based on its closeness to the *Tlr7* peak (within the 50% CI), its merge strength and RNA-seq in bone cells, we identified *Rhbdf2* as a likely causal gene associated with BV/TV and Tb.N. We therefore analyzed the bone phenotype of *Rhbdf2*^-/-^ mice, to validate the role of this gene in the modeling of the femoral cortex and trabeculae. *Rhbdf2* deletion affected all the examined trabecular traits as well as Ct.Th. While the effects on BV/TV and Tb.N were in line with the haplotype mapping, the *Rhbdf2* locus did not appear in any of the other traits. This however is expected, because the genetic architecture of the working cohort is such that the assumed contributing variant of *Rhbdf2* is diluted and compensated, resulting in a QTL detected only for the most affected traits. Noticeably, Tb.Sp differed greatly between the *Rhbdf2*^-/-^ and control mice but did not show up at the haplotype mapping; this might be due to (i) the great diversity of the wild-type mice in Tb.Sp, and/or (ii) the need for complete knockout rather than a mere SNP to detect significant changes in Tb.Sp, and/or (iii) the SNPs giving rise to *Trl*7 are functioning variants, with differential behavior affecting only BV/TV and Tb.N. A similar interpretation may be valid for the cortical phenotype of the *Rhbdf2*^-/-^ mice. Importantly, the significant QTL peak we found in our GWAS for BV/TV and Tb.N ended up revealing a gene that has an important skeletal function in both the trabecular and cortical bone compartments.

*Rhbdf2* encodes the iRhom2 protein, a polytopic membrane protein that is a catalytically inactive member of the rhomboid intramembrane serine proteases superfamily [59]. iRhom2 is necessary in macrophages for the maturation and release of the inflammatory cytokine tumor necrosis factor α (TNFα): it acts in the trafficking of TACE, the protease that releases active TNFα from its membrane-tethered precursor [60,61]. iRhom2 is also implicated in EGF-family growth factor signaling [62–64]. With a recent report of its role in trafficking of another protein, STING, it appears that iRhom2 may have a wider role in regulating membrane trafficking [65]. iRhom2 was also implicated in the regulation of CSF1R (macrophage stimulating factor 1 receptor), a critical regulator of osteoclasts differentiation and survival [61,66–68]. *In vivo*, *Rhbdf2* has been implicated in esophageal cancer, wound healing, bone marrow repopulation by monocytic cells, and inflammatory arthritis [63,69–71].

Further work will be needed to identify the mechanism by which iRhom2 controls bone homeostasis; a possible direction could involve a positive feedback loop that leads to differentiation of macrophages to osteoclasts. Indeed, iRhom2 stimulates the secretion of TNFα by macrophages [68,72]; hyperactivates EGFR [73,74]; and regulates CSF1R [75,76]. Although *Rhbdf2* is expressed in both the osteocyte and osteoclast lineages, one cannot rule out the possibility that this gene regulates bone remodeling by virtue of its expression in non-skeletal cells.

In summary, our analyses disclose several putative genes, several of which are newly linked to a role in bone biology. A confirmation of one such gene, *Rhbdf2*, provides the first conclusive evidence for its effects on bone microstructure. Importantly, this study is the first demonstration of a physiological role of *Rhbdf2*. This finding prompts future investigations to elucidate the exact mechanism of action of *Rhbdf2* and its contribution to osteoporosis in humans.

## Materials and Methods

### Mice

Mice aged 10 to 13 weeks (male *n =* 103; female *n* = 71), from 34 different CC lines (average of 5 mice per line) were used in this study. The mice were at inbreeding generations of 11 to 37, which correspond to 80-99.9% genetic homozygosity, respectively. The mice were bred and maintained at the small animal facility of the Sackler Faculty of Medicine, Tel Aviv University (TAU), Israel. They were housed on hardwood chip bedding in open-top cages, with food and distilled water available *adlibitum*, in an identical controlled environment (temperature = 25 ± 2°C; 60% ≤ humidity ≤ 85%) and a 12-hour light/dark cycle. All experiments protocols were approved by the Institutional Animal Care and Use Committee (IACUC M-13-014) at TAU, which follows the NIH/USA animal care and use protocols. The *Rhbdf2* knock out mice and their WT counterparts were bred and maintained at the University of Oxford as approved by license PPL80/2584 of the UK Home Office.

### Specimen collection

Mice were intraperitoneally euthanized with cervical dislocation performed approximately one minute after breathing stops owing to 5% Isoflurane inhalation. The *Rhbdf2* knock out mice and their WT counterparts were euthanized by inhalation of a rising concentration of carbon dioxide followed by dislocation of the neck. Left femora were harvested and fixed for 24 hours in 4% paraformaldehyde solution, and then stored in 70% ethanol.

### μCT evaluation

Whole left femora from each mouse were examined as described previously [77] by a [μCT system (μCT 50, Scanco Medical AG, Switzerland). Briefly, scans were performed at a 10-μm resolution in all three spatial dimensions. The mineralized tissues were differentially segmented by a global thresholding procedure [78]. All morphometric parameters were determined by a direct 3D approach [79]. Parameters analyzed were determined in the metaphyseal trabecular bone, which included trabecular bone volume fraction (BV/TV; %), trabecular thickness (Tb.Th; μm), trabecular number (Tb.N; mm^−1^), trabecular connectivity density (Conn.D; mm^−3^), trabecular structure model index (SMI), and trabecular separation (Tb.Sp; mm). Two additional parameters are characteristics of the mid-shaft diaphysis section, and include volumetric bone mineral density (vBMD; mgHA/cm^3^ [mg Hydroxy-Apatite per cm^3^]) and cortical thickness (Ct.Th; mm). All parameters were generated and termed according to the Guidelines for assessment of bone microstructure in rodents using micro–computed tomography [80].

### Genotvping

A representative male mouse from each line was initially genotyped with a high mouse diversity array (MDA), which consists of 620,000 SNPs (Durrant et al., 2011). After about two intervals of 4 generations of inbreeding, all the CC lines were regenotyped by mouse universal genotype array (MUGA, 7,500 markers) and finally with the MegaMuga (77,800 markers) SNP array to confirm their genotype status [19]. The founder-based mosaic of each CC line was reconstructed using a hidden Markov model (HMM) in which the hidden states are the founder haplotypes and the observed states are the CC lines, to produce a probability matrix of descent from each founder. This matrix was then pruned to about 11,000 SNPs by averaging across a window of 20 consecutive markers for faster analyses and reduction of genotyping errors [81].

### Statistical analyses and data acquisition

All statistical analyses were performed with the statistical software R (R core development team 2009), including the package happy.hbrem [82].

#### Heritability and covariate effects

Broad-sense heritability (H^2^) was obtained for each trait by fitting the trait (the independent variable) to the CC line label in a linear regression model that incorporates relevant covariates (sex, age, batch, month, season, year, and experimenter). ANOVA test was used to compare a null model (in which all dependent variables are set to 0) with linear models that fit the covariates and the CC line labels to the examined trait. Practically, the difference between the residual sum of squares (RSS; 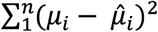) of the covariates model and that of the CC-line labels can be seen as the net genetic contribution to the trait. Thus, this difference divided by that of the covariate model gives an estimation of the heritability. Each covariate was calculated separately, by dividing the RSS difference between the null and full model with that of the null model. Let *F*_0_ be the model that fits the trait to the covariates; *F*_1_ the model that fits the trait to the covariates and the CC line label; and *F*_00_ the null model. Then, employing ANOVA, heritability is:

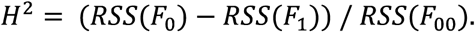

Similarly, the effects for each covariate were computed separately, by fitting each in *F*_0_. The covariate effect is thus:

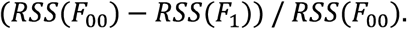

H^2^n was derived from H^2^ according to Atamni *et al* [23].

#### Haplotype mapping

Each trait was fitted in a multiple linear regression model to the probability matrix of descent from each founder, including sex and age as covariates. The expected trait value from two ancestors, termed the genetic fit, is:

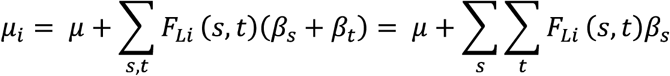

where *μ* is a normally distributed trait mean, with sex and age incorporated; *F_Li_* (*s, t*) is the probability of descent from founders *s* and *t;* and *β_s_* + *β_t_* is the additive effect of founders *s* and *t*. Because ∑*c_s_*∑*_t_ F_Li_* (*s, t*) = 2 for a diploid organism, the maximum likelihood estimates *β̂_s_* are not independent. Thus, they are expressed here as differences from the WSB/EiJ founder effect, so that *β̂_WSB_* = 0. Number of members per line was weighted and integrated in the linear model. ANOVA was then used to compare this model with a null model where the founder effects are all set to 0; the resulting F-statistic yielded the significance of the genetic model vs. the null model and the negative 10-base logarithms of the P values (logP) were recorded.

Permutations of the CC lines between the phenotypes were used to set significance thresholds levels. Founder effects are the estimates derived from the multiple linear regression fit above.

Regional heritability (H_r_^2^) was hereafter computed by ANOVA as in the broad-sense heritability computation, except that here null linear regression fit was compared with a genetic linear regression fit with the probability matrix of the founder descent at the peak QTL as the explanatory variable.

False discovery rate (FDR) was calculated using the p.adjust function in R, with the method “BH” (Benjamini-Hochberg [83]).

#### Confidence intervals

Confidence intervals (CIs) were obtained both by simulations and by the quick method of Li, 2011 [84], In the simulations, we resampled the residuals of the original linear regression fit at the peak of each QTL and rescanned 100 intervals within 7-10 Mb of the original loci to find the highest logP. Accordingly, following Durrant *et al.* [22], 1000 QTLs were simulated: if *t̂_i_* is a random permutation of the residuals of fitted genetic model at the QTL peak, and *K* is a marker interval in a neighborhood of 3.5 to 5 Mb of the QTL peak *L*, a set of values for each trait, *Z_iK_* is provided by:

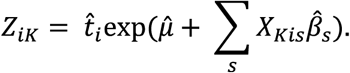

#### Merge analysis

In the merge analysis the eight founder strains are partitioned and merged according to the strain distribution pattern (SDP) of the alleles at the quantitative trait nucleotides (QTN) within a given QTL (formerly obtained by the initial mapping). If we denote the polymorphism as *p*, then *X_p_ =* 1 if *s* has allele *a* at *p*, and *X_p_ =* 0 otherwise [24], Then, at *p*, the probability of *i* to inherit alleles *a and b* from *s and t, respectively*, within *L* is

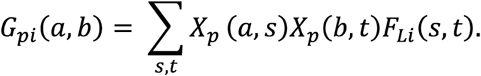

This merges the founder strains by *p.* The expected trait value in the merged strains can now be inferred by

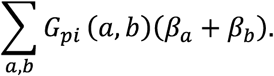

Because this is a sub-model of the QTL model, it is expected to yield higher logP values due to a reduction in the degrees of freedom. Significance was obtained by comparing the merge model with the QTL model. Individual genes were extracted from the Sanger mouse SNP repository (http://www.sanger.ac.uk/sanger/Mouse_SnpViewer).

#### Merge strength

We ranked the list of genes under each QTL according to the density of merge logPs associated with them: only genes that had merge logPs above the haplotype mapping reading, and above the threshold, plus logP=1 were included. We then computed the relative density according to the density of a given gene’s merge logPs versus the locus’ merge logP density. Let *g* be the region encompassed by a gene; *l* the region encompassed by a QTL; and *mp* the merge logP values above the haplotype P values plus 1. Then *g_i_*(*mp*) = 1 if at SNP *i* there exists a *mp* and 0 otherwise. Similarly *l_i_*(*mp*) = 1 if at SNP *j* there exists a *mp* and 0 otherwise. The merge strength (*M_strength_*) is therefore:

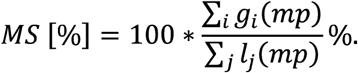

#### RNA-seq data

RNA-seq data from osteoclasts and osteocytes was obtained from gene expression omnibus (GEO) database (accession numbers GSE72846 and GSE54784) and mapped to the *mus musculus* assembly mm10 using tophat v. 2 [85], Read counts were then casted on the loci of interest using the R (R Core Team 2015) package GenomicAlignments and raw read counts were taken. For the osteocytes, the data of basal level day 3 was averaged.

MS score. Based on the merge analysis and RNA-seq data we ranked each gene according to the score in each category: *MS(i)* = *ln(M_strength_(i))* + (*S_RNA-seq1_(i)* + *S_RNA-seq2_(i)*), where *MS*, *M_strength_*, *S_RNA-seq1_*, and *S_RNA-seq2_* refer to the merge and sequencing score, merge rank (or strength), osteoclast RNA-seq score, and osteocyte RNA-seq score of a given gene (*i*), respectively. *S_RNA-seq1,2_* scores were generated according to the formula 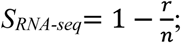 where *r* is the local gene rank (sorted by the raw read count) and n is the total number of genes at the locus, where *n*=10, if *n*>10.

## Acknowledgments

This study was supported by Tel Aviv University starter funds and by Israel Science Foundation (ISF) grant 1822/12 to YG, by Wellcome Trust grants 085906/Z/08/Z, 075491/Z/04, and 090532/Z/09/Z to RM, core funding by Tel-Aviv University to FI, Wellcome Trust grant 101035/Z/13/Z and the Medical Research Council (programme number U105178780) to MF, and by a fellowship from the Edmond J. Safra Center for Bioinformatics at Tel-Aviv University to RL.

## Supporting information captions

**Figure S1** *Trait distributions for Males and Females.* **A1**, **A2**: Trabecular and cortical bone, respectively, males. **B1, B2:** trabecular and cortical bone, respectively, females. X-axis is the lines, y-axis is the trait means A. From top left, counter-clockwise: BV/TV (%), Tb.N (mm-1), Tb.Th (um), Conn.D (mm-3), SMI, and Tb.Sp (mm). **B:** Left, vBMD (mgHA/cm3); right, Ct.Th (mm). Color codes group line(s) which significantly differ from other groups. Lines are ordered inconsistently among the traits, according to trait-specific descending order.

**Figure S2** *Haplotype association maps for the trabecular and cortical traits.* X-axis is the position on the chromosome, y-axis is the −logP value of the association. Lower threshold represents the 95th percentile of 200 simulations, and top represents the 9th percentile. Loci above the 99% cut-off were further investigated. From top to bottom: Conn.D, SMI.

**Figure S3** *Confidence interval simulations.* Loci at a neighborhood of 3-5 Mb around the original locus were simulated by permuting the residual sum of squares of the related phenotype. Maximum logP was obtained along with its relative position in Mb to the original QTL (histograms, left panels), and with the number of markers from the original QTL (boxplots, right panels). (*A)* and (*B*) show simulations results for the BV/TV and Tb.N loci. These determined with high confidence that the peak QTL is responsible for the effect seen in the haplotype scan, thus the narrow CI; (*C)* to (*F)* show simulation results for Tb.Th, Tb.Sp, Ct.Th, and vBMD, respectively; note the narrow CI for *Crl2* (vBMD), wide for *Trl8* (Tb.Th), and wider still for *Crl1(*Ct.Th).

**Figure S4** *RNA-seq of osteoclasts.* Gene names are on the right of each plot. Green represents plus-stranded genes, black represents minus-stranded genes. Y-axis is the raw expression count, where the negative scale refers to minus-stranded gene count. Each bracket corresponds to a particular gene; left-to-right green (black) brackets fit green (black) top-to-bottom gene-names.

**Figure S5** *RNA-seq of osteocytes.* Gene names are on the right of each plot. Orange represents plus-stranded genes, black represents minus-stranded genes. Y-axis is the raw expression count, where the negative scale refers to minus-stranded gene count. Each bracket corresponds to a particular gene; left-to-right orange (black) brackets fit orange (black) top-to-bottom gene names

**Table SI** *Pearson’s correlations between measured μCTtraits.* Pairwise correlations for each trait are given as Pearson’s *r.* In bold are the 5 most strong correlations.

**Table S2** *Covariates effects. LogP is the negative logarithm of P value.* Effects of covariates (i.e., the degree to which each covariate explains the phenotypic difference). Values were determined by regressing the covariates along with the CC lines, and running an ANOVA test. Note the covariates prominent effect on the cortical traits. The batch effect was the strongest, affecting Tb.Th, Ct.Th, and vBMD.

**Table S3** *Trait values per CC line included in this study (for both sexes combined).* Trait means for each line, including number of members and standard error (SE). NA means data were not available.

**Table S4** *A comprehensive list of all genes under the 95% confidence-interval (CI) for the QTLs identified in this study.* Light blue is the 95% CI region; blue is the 90% CI region; and pink is the 50% CI region. Merge strength column gives the normalized dosage of merge logP values as outlined in the methods section. Gene symbols in bold type are discussed in the manuscript. Genes under the QTLs for the cortical and trabecular traits. Light blue is the 95% CI, blue is the 90% CI, and red is the 50% CI. “Merge Strength” refers to the proportion of merge logP values constricted to the region of the specific gene. Note that for Trl7 which is common to BV/TV and Tb.N, the average values between the two are provided.

